# Mechanophysiology of endometriosis: a non-dimensional physiomarker to detect retrograde flow

**DOI:** 10.1101/2024.05.13.593987

**Authors:** Guy Elisha, Neelesh A. Patankar

**Affiliations:** Department of Mechanical Engineering Northwestern University, Evanston, IL, USA; Department of Engineering Sciences and Applied Mathematics, Northwestern University, Evanston, IL, USA

**Keywords:** endometriosis, uterus, fallopian tube, peristalsis, physiomarker

## Abstract

Endometriosis affects a significant portion of fertile-age women, often leading to infertility and a substantial decline in quality of life. Despite its prevalence, current diagnostic methods are limited, focusing on assessing the presence or absence of endometrial lesion, rather than the origin of the disorder. Thus, resulting in underdiagnosis. A potential mechanics-based metric for diagnosing endometriosis is proposed here by leveraging the retrograde menstruation hypothesis. By examining the interplay between uterine and fallopian tube peristalses, a non-dimensional physiomarker is introduced to signify the onset of retrograde flow. The analysis reveals that increased uterine contractile activity, coupled with decreased fallopian tube contractile activity, correlates with retrograde flow, suggesting a predisposition to endometriosis. This mechanophysiology-based approach offers a promising avenue for origin based diagnosis, with the proposed non-dimensional physiomarker – the endometriosis number – serving as a potential indicator of endometrial cell migration and the onset of endometriosis.

## 1 Introduction

Endometriosis is estimated to affect roughly 10% of fertile-age women, accounting for up to 50% of female infertility cases, and can have a debilitating effect on women’s lives [1, 2]. The severe pain interferes with daily activities, relationships, and livelihood [3]. The disorder is characterized by the endometrium tissue, which is normally inside the uterus, growing outside the uterine cavity. During healthy menstrual cycle, the endometrium tissue inside the uterus thickens, breaks down, and is shed as menstrual blood. During endometriosis, the same process occurs, but the tissue also grows in places outside the uterus covering the ovaries and fallopian tubes. Consequently, after shedding, the blood cannot exit the body [4].

Despite its prevalence, endometriosis is under-researched and it often goes undiagnosed [5, 6]. Accurate diagnosis of endometriosis requires invasive techniques, contributing to a lack of diagnosis [3, 7]. In recent years, there have been repeated calls for finding more efficient diagnostic approaches [3, 7]. Agarwal et al.[3] proposed that diagnosing endometriosis should prioritize symptoms and their origins over assessing the presence or absence of endometrial lesions. However, there is no procedure to quantify or measure the origin of endometriosis. In this work, we explore a potential mechanics-based metric to diagnose endometriosis targeting the origin.

Retrograde menstruation hypothesis is a widely accepted plausible explanation for the development of endometriosis [8, 9]. It suggests that during menstruation, some menstrual blood containing endometrial cells flows backward through the fallopian tubes into the pelvic cavity instead of being expelled from the body. These endometrial cells then implant and grow in various pelvic locations, leading to the development of endometriosis.

Here we focus on finding a diagnostic metric for endometriosis motivated by the success of mechanics-based tools and criteria to diagnose and assess disease progression in other organs [11, 12]. Mechanics-based approaches to investigate the physiology (mechanophysiology) of both the uterus and the fallopian tubes have been reported in the past [13, 14, 15, 16, 17, 18, 19]. Studies so far have treated the uterus and fallopian tubes as separate entities, without exploring potential emergent outcomes due to their interconnected influence. Here, we explore if the competition between uterine peristalsis and fallopian tubes’ peristalsis gives insights into when flow becomes retrograde (from the cervix to the fundus into the fallopian tubes, see Fig. 1). By doing so, we come up with a non-dimensional physiomarker (a physics-based metric) which marks the critical condition for the onset of retrograde flow.

**Figure 1:**
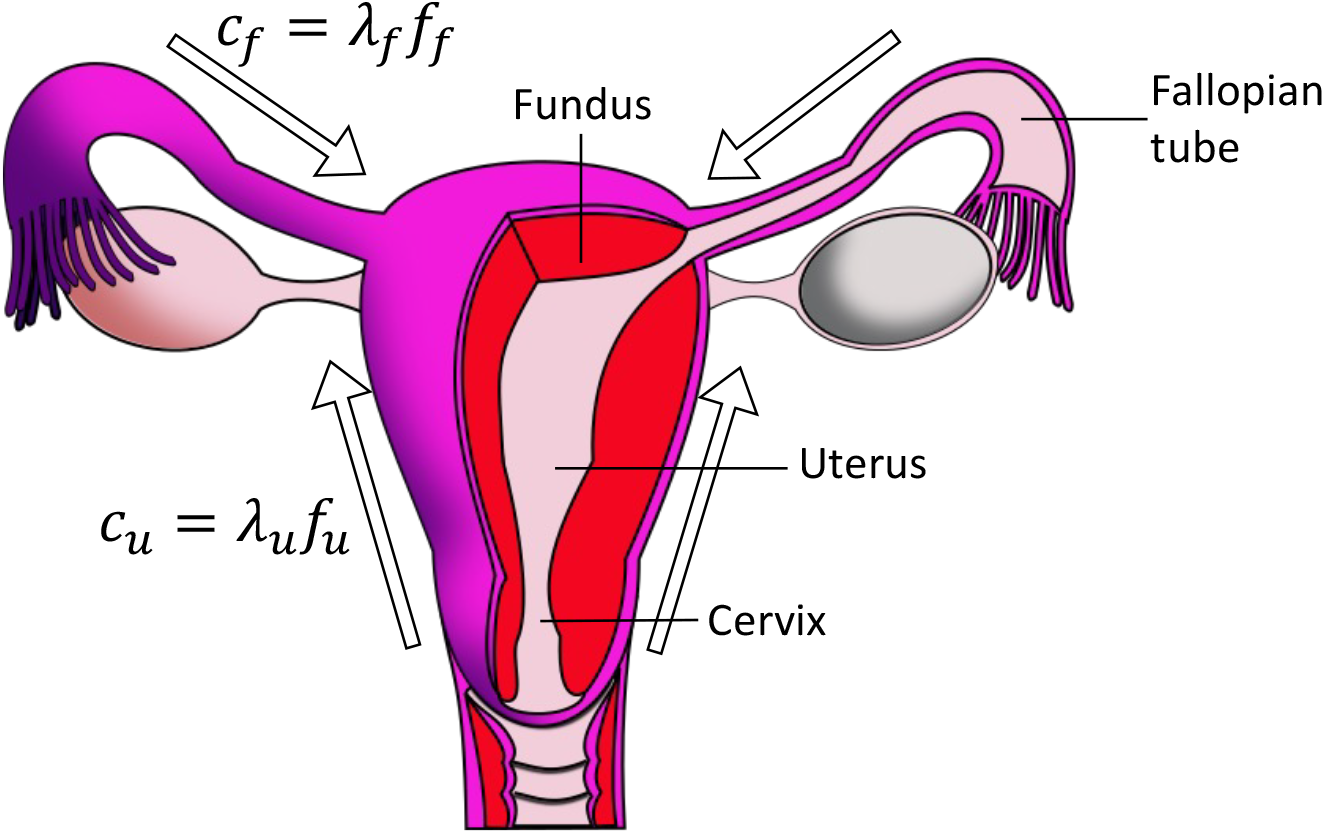
Uterine cavity and the fallopian tubes intersect around the fundus. The directions of the muscle contractions waves of the uterus and the fallopian tubes are marked with arrows, both directing towards the fundus during potential retrograde flow. Images reproduced with permission from [10]. Labels and arrows were added to the original image.

## 2 Mathematical analysis

### 2.1 Uterus

Let *a*_*u*_ represent the mean half-height of the uterus channel and *λ*_*u*_ denote the uterus peristaltic wavelength. According to Eytan et al. [13] and Aranda et al. [20], it holds that *a*_*u*_*/λ*_*u*_ *<<* 1. Additionally, uterine Reynolds number (*Re*_*u*_) satisfies

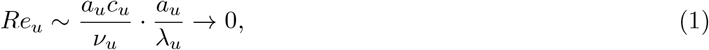

where *c*_*u*_ is the uterine peristaltic wave speed and *ν*_*u*_ denotes kinematic viscosity of the uterine fluid. Thus, uterine flow is almost inertialess. Given these assumptions, we utilize the solution for a peristaltic flow inside a two-dimensional planar channel geometry derived by Shapiro et al. [21] given by

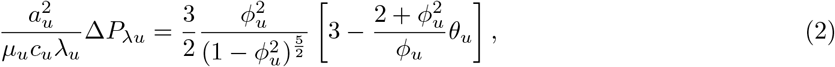

where *μ*_*u*_ represents the dynamic viscosity of the uterine fluid, Δ*P*_*λu*_ denotes the uterine pressure rise per wavelength, and *ϕ*_*u*_ = *b*_*u*_*/a*_*u*_ indicates the uterine amplitude ratio, where *b*_*u*_ stands for the half-amplitude of the uterine peristaltic wave. Additionally, 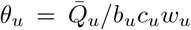 signifies the dimensionless time-averaged flowrate in the uterus, where 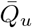 represents the time-averaged volumetric flowrate at each cross-section along the uterus, and *w*_*u*_ denotes the width of the uterine channel [21]. The pressure rise over the entire uterus length is expressed by

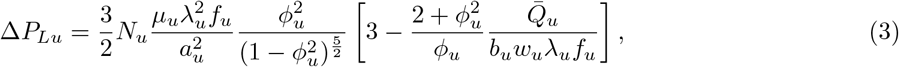

where Δ*P*_*Lu*_ signifies the pressure at the junction of the uterus and the fallopian tubes (the fundus), *f*_*u*_ is the uterus peristaltic frequency, and *N*_*u*_ indicates the number of wavelengths in the uterine length. Note that we assume that the pressure at the entrance of the uterus (the cervix) is zero.

### 2.2 Fallopian Tubes

Similarly, to express the pressure rise over the length of a fallopian tube (Δ*P*_*Lf*_ ), we again assume that *a*_*f*_ */λ*_*f*_ *<<* 1 and *Re →* 0 (*a*_*f*_ and *λ*_*f*_ denote the radius of the fallopian tube and the fallopian tube peristaltic wavelength, respectively). It follows from Shapiro et al. [21] that

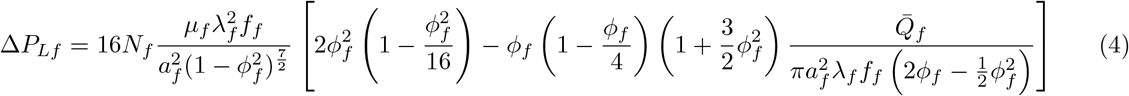

where *μ*_*f*_ represents the dynamic viscosity of the fallopian tube fluid, *f*_*f*_ is the fallopian tube peristaltic frequency, *N*_*f*_ indicates the number of wavelengths in the length of the fallopian tube, 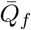 signifies the time-averaged volumetric flowrate at each cross-section along the fallopian tube, *ϕ*_*f*_ = *b*_*f*_ */a*_*f*_ indicates the fallopian tube amplitude ratio, and *b*_*f*_ stands for the half-amplitude of the fallopian tube peristaltic wave. Note that Eq. (4) represents the flow through one fallopian tube, but the system consists of two tubes intersecting with the uterus. Furthermore, by convention, the traveling contraction in the fallopian tube is in the opposite direction to the one of the uterus (Fig. 1). Hence, the net flow in the fallopian tubes from the fundus toward the ovaries is given by 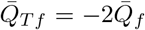. Finally, it is assumed that the pressure at the ovarian end of the fallopian tubes (in the pelvic cavity) is zero.

### 2.3 Critical Condition

At the junction of the uterus and the fallopian tubes, it is required that Δ*P*_*Lu*_ = Δ*P*_*Lf*_ and 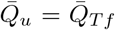 to satisfy steady-state momentum and mass conservation equations. These conditions when imposed in Eqs. (3) and (4) give the solutions for the volumetric flow rate and pressure drop as depicted in Fig. 2a. If the solution for the volumetric flow rate 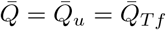 is such that 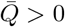, then the flow is retrograde; there is no retrograde flow if 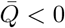.

**Figure 2:**
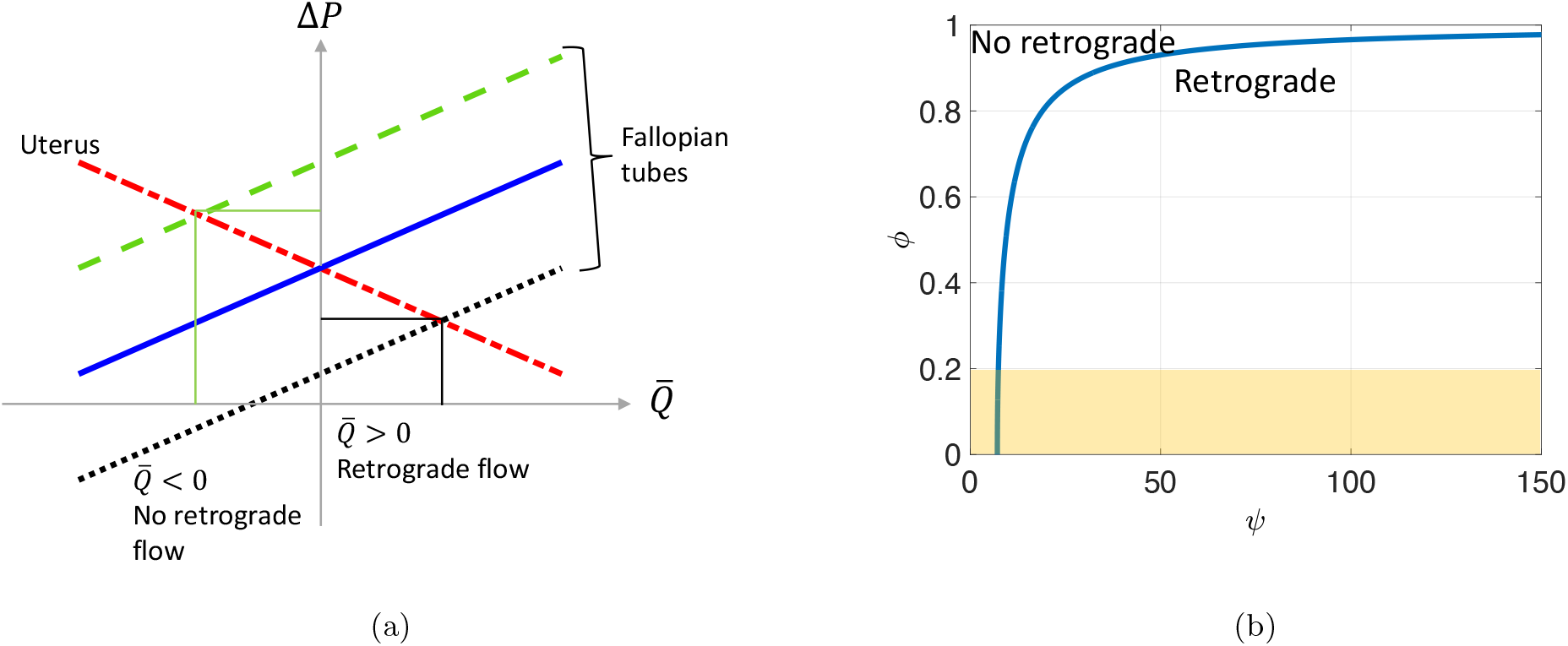
(a) A sketch of lines representing the pressure rise over the length of the uterus (Δ*P*_*Lu*_) and the fallopian tubes (Δ*P*_*Lf*_ ) as a function of their respective time-averaged volumetric flowrates (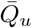 and 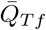). The intersection of the two lines, for specific parameters in Eqs. (3) and (4), gives the solution for the volumetric flowrate 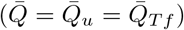 and the pressure drop (Δ*P* = Δ*P*_*Lu*_ = Δ*P*_*Lf*_ ). If the solution is such that 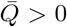 (e.g. red and black lines) then the flow is retrograde. Alternately, if 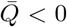 (e.g. red and green lines) then there is no retrograde flow. (b) Graph showing the dimensionless endometriosis number *ψ* vs. *ϕ*, derived from Eq. (7), along with possible range for clinical values for *ψ* and *ϕ* (orange rectangle). The graph separates the *ϕ*–*ψ* space into retrograde and no retrograde scenarios.

We can establish a critical condition for retrograde flow using Fig 2a. Consider parameters such that the uterine peristaltic flow is represented by the red line (Eq. 3) in Fig 2a. Now consider three different scenarios for fallopian tube peristaltic flows represented by the green, blue, and black lines (Eq. 4) in Fig 2a. It is seen that if the fallopian tube peristalsis is represented by the black line then the flow will be retrograde because it intersects the red uterine flow line for 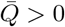, whereas if the fallopian tube peristalsis is represented by the green line then the flow won’t be retrograde (Fig 2a). It is evident that the blue line case is the borderline scenario between retrograde and no retrograde flow because 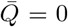 when it intersects with the red uterine peristalsis line. This shows that, *no retrograde* flow will take place if

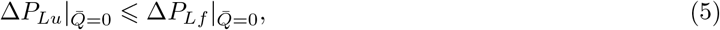

where the equality represents the critical condition. We substitute Eqs. (3) and (4) into Eq. (5) to derive

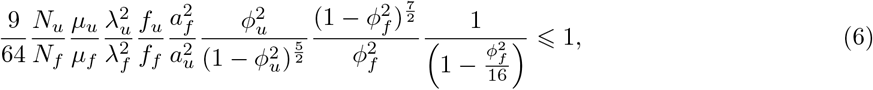

for no retrograde flow. We define a non-dimensional number 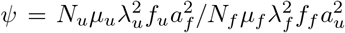 and assume that *ϕ*_*u*_ *≈ ϕ*_*f*_ (= *ϕ*). Thus, the no retrograde flow condition can be expressed in terms of parameter *ψ*, such that

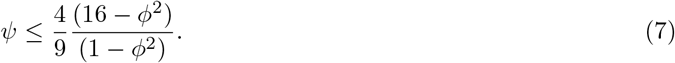

By setting up this inequality, we find an *endometriosis number ψ* as a potential *physiomarker*, which is based on measurable physical quantities on a patient specific basis. It can be used to quantify the critical condition for the onset of retrograde flow potentially causing endometriosis. The plot of the equation above is displayed in Fig. 2b and is discussed in greater details in the following section.

Notice that

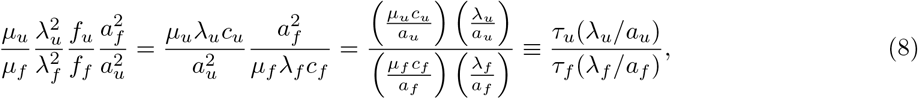

where *τ*_*u*_ and *τ*_*f*_ are the scales of viscous shear stresses on the walls of the uterus and the fallopian tubes, respectively. To ensure that there is no retrograde flow, the fallopian tubes’ viscous resistance must be greater than that of the uterus.

## 3 Discussion

The graph plotted in Fig. 2b, obtained from Eq. (7), delineates a space wherein distinct scenarios of retrograde flow and no retrograde flow can be distinguished. Thus, by extracting physical measurements of a particular individual, such as contraction frequency and tube thickness (referenced in Table 1), we can compute *ψ* and *ϕ*, and thereby determine the presence or absence of retrograde flow.

**Table 1:** List parameters and their values gathered from prior clinical studies.

| Parameter | Definision | Uterus | Fallopian Tube | Units | References |
| --- | --- | --- | --- | --- | --- |
| 2a | Mean height of the uterus channel or diameter of the fallopian tubes | 1-4 | 1-4 | mm | [22, 13, 23, 24, 18, 25] |
| 2b | Amplitude of the peristaltic wave | 0.05-0.2 | 0.03-0.64 | mm | [23, 22, 26] |
| c | Wave speed | 0.5-1.9 | 1-3 | mm/s | [22, 27, 28] |
| f | Peristaltic frequency | 0.03-0.08 | 0.025-0.133 | 1/s | [23, 22, 29, 30] |
| N | Number of wavelengths | 1.5-3.8 | 2-4<sup>1</sup> |  | [31, 32] |
| $\lambda$ | Contraction wavelength | 10-30 | 10-50<sup>2</sup> | mm | [22, 28, 29] |

1.Calculated using the length and wavelength of the fallopian tubes.
2.Calculated using the wavelength and wave speed of the fallopian tubes.

We gather the necessary parameter values for calculating *ψ* and *ϕ* from prior clinical studies, detailed in Table 1, and make the assumption that *μ*_*u*_ = *μ*_*f*_ . We find that the clinically plausible range for the endometriosis number *ψ* is approximately within 0.01 to 150. Smaller *ψ* values (ranging between 0 to 50) are far more common. Moreover, empirical data indicates that *ϕ →* 0 since *a >> b*. Hence, Eq. (7) simplifies to *ψ >* 7.1, indicating that retrograde flow manifests when *ψ >* 7.1. This conclusion remains consistent when examining various combinations of *ψ* and *ϕ* plotted in Fig. 2b. Importantly, we deduce that retrograde flow presence is solely dictated by the endometriosis number *ψ*.

Leyendecker et al. [33] reported significant increase in the peristaltic activity (frequency) of the uterus in infertile female diagnosed with endometriosis. Kunz and Leyendecker [34] and Kissler et. al [35] observed hypercontractile activity of the uterus in certain patients, noting that this condition may be involved in the development of endometriosis. Xia et al. [29] have recorded an opposite trend in the fallopian tube. They noticed that the fallopian tubes’ contraction frequency of controls was higher than the ones of patients with endometriosis. These findings align with our results, as the endometriosis number *ψ* and the critical condition for retrograde flow depend on uterine and fallopian tube contraction activity. Our analysis shows that an increase in the uterine contractile activity and a decrease in the fallopian tubes’ contractile activity favor retrograde flow.

To compute the patient-specific value of the endometriosis number, measurements from both the uterus and fallopian tubes are required. Uterine thickness (2*a*_*u*_) and contraction properties (*λ*_*u*_, *c*_*u*_, *f*_*u*_, *N*_*u*_) can be assessed using transvaginal ultrasound [23]. However, there is currently no established method for obtaining comparable data (*a*_*f*_, *λ*_*f*_, *c*_*f*_, *f*_*f*_, *N*_*f*_ ) for the fallopian tubes in vivo. The parametric values for the fallopian tubes listed in Table 1 have been gathered in vitro [29, 25]. If similar non-invasive techniques can be developed to obtain fallopian tube data, as has been done for the uterus, our proposed approach could offer a less-invasive alternative to laparoscopy. Laparoscopy, a surgical procedure currently considered the gold standard for endometriosis diagnosis, is invasive in nature [3, 7].

This study provides a mechanophysiology-based analysis aimed at diagnosing an individual’s predisposition to developing endometriosis. By computing the endometriosis number *ψ* introduced in this work, we can determine the presence of retrograde flow, indicating a likelihood of endometrial cell migration beyond the uterine cavity and suggesting the onset of endometriosis. Future work focused on testing the endometriosis number *ψ* though clinical measurements of both patients and controls is recommended.

## Acknowledgments

This work was funded by the by the National Science Foundation (OAC grant 1931372).

## References

[1] Giudice, L. C., 2010, “Endometriosis,” New England Journal of Medicine, 362(25), pp. 2389–2398.

[2] Carter, J. E., 1994, “Combined hysteroscopic and laparoscopic findings in patients with chronic pelvic pain,” The Journal of the American Association of Gynecologic Laparoscopists, 2(1), pp. 43–47.

[3] Agarwal, S. K., Chapron, C., Giudice, L. C., Laufer, M. R., Leyland, N., Missmer, S. A., Singh, S. S., and Taylor, H. S., 2019, “Clinical diagnosis of endometriosis: a call to action,” American journal of obstetrics and gynecology, 220(4), pp. 354–e1.

[4] Horne, A. W. and Missmer, S. A., 2022, “Pathophysiology, diagnosis, and management of endometriosis,” bmj, 379.

[5] Horne, A. W., Saunders, P. T., Abokhrais, I. M., and Hogg, L., 2017, “Top ten endometriosis research priorities in the UK and Ireland,” The Lancet, 389(10085), pp. 2191–2192.

[6] Seear, K., 2016, The makings of a modern epidemic: endometriosis, gender and politics, Routledge.

[7] Lin, Y.-H., Chen, Y.-H., Chang, H.-Y., Au, H.-K., Tzeng, C.-R., and Huang, Y.-H., 2018, “Chronic niche inflammation in endometriosis-associated infertility: current understanding and future therapeutic strategies,” International journal of molecular sciences, 19(8), p. 2385.

[8] Hill, C. J., Fakhreldin, M., Maclean, A., Dobson, L., Nancarrow, L., Bradfield, A., Choi, F., Daley, D., Tempest, N., and Hapangama, D. K., 2020, “Endometriosis and the fallopian tubes: Theories of origin and clinical implications,” Journal of clinical medicine, 9(6), p. 1905.

[9] Bricou, A., Batt, R. E., and Chapron, C., 2008, “Peritoneal fluid flow influences anatomical distribution of endometriotic lesions: why Sampson seems to be right,” European Journal of Obstetrics & Gynecology and Reproductive Biology, 138(2), pp. 127–134.

[10] Wikimedia Commons, 2024, “Uterus,” https://commons.wikimedia.org/wiki/File:Uterus.png

[11] Halder, S., Pandolfino, J. E., Kahrilas, P. J., Koop, A., Schauer, J., Araujo, I. K., Elisha, G., Kou, W., Patankar, N. A., and Carlson, D. A., 2023, “Assessing mechanical function of peristalsis with functional lumen imaging probe panometry: Contraction power and displaced volume,” Neurogastroenterology & Motility, 35(12), p. e14692.

[12] Zhao, T. Y., Johnson, E. M., Elisha, G., Halder, S., Smith, B. C., Allen, B. D., Markl, M., and Patankar, N. A., 2023, “Blood–wall fluttering instability as a physiomarker of the progression of thoracic aortic aneurysms,” Nature Biomedical Engineering, 7(12), pp. 1614–1626.

[13] Eytan, O. and Elad, D., 1999, “Analysis of intra-uterine fluid motion induced by uterine contractions,” Bulletin of Mathematical Biology, 61(2), pp. 221–238.

[14] Eytan, O., Jaffa, A. J., and Elad, D., 2001, “Peristaltic flow in a tapered channel: application to embryo transport within the uterine cavity,” Medical engineering & physics, 23(7), pp. 475–484.

[15] Aranda, V., Cortez, R., and Fauci, L., 2011, “Stokesian peristaltic pumping in a three-dimensional tube with a phase-shifted asymmetry,” Physics of fluids, 23(8).

[16] Yaniv, S., Jaffa, A. J., and Elad, D., 2012, “Modeling embryo transfer into a closed uterine cavity,”.

[17] Aranda, V., Cortez, R., and Fauci, L., 2015, “A model of Stokesian peristalsis and vesicle transport in a three-dimensional closed cavity,” Journal of biomechanics, 48(9), pp. 1631–1638.

[18] Ashraf, H., Siddiqui, A., and Rana, M., 2018, “Analysis of the peristaltic-ciliary flow of Johnson– Segalman fluid induced by peristalsis-cilia of the human fallopian tube,” Mathematical biosciences, 300, pp. 64–75.

[19] Salman, M. R., Asghar, Z., Javid, K., Waqas, M., et al., 2019, “Fallopian tube analysis of the peristaltic-ciliary flow of third grade fluid in a finite narrow tube,” Journal of Physics: Conference Series, Vol. 1362, p. 012157.

[20] Aranda, V., Cortez, R., and Fauci, L., 2011, “Stokesian peristaltic pumping in a three-dimensional tube with a phase-shifted asymmetry,” Physics of fluids, 23(8), p. 081901.

[21] Shapiro, A. H., Jaffrin, M. Y., and Weinberg, S. L., 1969, “Peristaltic pumping with long wavelengths at low Reynolds number,” Journal of fluid mechanics, 37(4), pp. 799–825.

[22] Eytan, O., Jaffa, A. J., Har-Toov, J., Dalach, E., and Elad, D., 1999, “Dynamics of the intrauterine fluid–wall interface,” Annals of biomedical engineering, 27, pp. 372–379.

[23] Meirzon, D., Jaffa, A., Gordon, Z., and Elad, D., 2011, “A new method for analysis of non-pregnant uterine peristalsis using transvaginal ultrasound,” Ultrasound in obstetrics & gynecology, 38(2), pp. 217–224.

[24] Giri, S., Nayak, B., and Mohapatra, J., 2021, “Thickened endometrium: when to intervene? A clinical conundrum,” The Journal of Obstetrics and Gynecology of India, 71(3), pp. 216–225.

[25] Guan, J. and Watrelot, A., 2019, “Fallopian tube subtle pathology,” Best Practice & Research Clinical Obstetrics & Gynaecology, 59, pp. 25–40.

[26] Nadasy, G., Laszlo, A., Monos, E., and Zsolnai, B., 1988, “Spontaneous periodic contraction of the ampullar segment of the human fallopian tube in vitro.” Acta Physiologica Hungarica, 72(1), pp. 13–21.

[27] Dodds, K. N., Staikopoulos, V., and Beckett, E. A., 2015, “Uterine contractility in the nonpregnant mouse: changes during the estrous cycle and effects of chloride channel blockade,” Biology of reproduction, 92(6), pp. 141–1.

[28] Talo, A. and Pulkkinen, M., 1982, “Electrical activity in the human oviduct during the menstrual cycle,” American Journal of Obstetrics and Gynecology, 142(2), pp. 135–147.

[29] Xia, W., Zhang, D., Ouyang, J., Liang, Y., Zhang, H., Huang, Z., Liang, G., Zhu, Q., Guan, X., and Zhang, J., 2018, “Effects of pelvic endometriosis and adenomyosis on ciliary beat frequency and muscular contractions in the human fallopian tube,” Reproductive Biology and Endocrinology, 16, pp. 1–7.

[30] Wånggren, K., Stavreus-Evers, A., Olsson, C., Andersson, E., and Gemzell-Danielsson, K., 2008, “Regulation of muscular contractions in the human Fallopian tube through prostaglandins and progestagens,” Human Reproduction, 23(10), pp. 2359–2368.

[31] Lyons, E. A., Taylor, P. J., Zheng, X. H., Ballard, G., Levi, C. S., and Kredentser, J. V., 1991, “Characterization of subendometrial myometrial contractions throughout the menstrual cycle in normal fertile women,” Fertility and sterility, 55(4), pp. 771–774.

[32] Orisaka, M., Kurokawa, T., Shukunami, K.-I., Orisaka, S., Fukuda, M. T., Shinagawa, A., Fukuda, S., Ihara, N., Yamada, H., Itoh, H., et al., 2007, “A comparison of uterine peristalsis in women with normal uteri and uterine leiomyoma by cine magnetic resonance imaging,” European Journal of Obstetrics & Gynecology and Reproductive Biology, 135(1), pp. 111–115.

[33] Leyendecker, G., Kunz, G., Wildt, L., Beil, D., and Deininger, H., 1996, “Uterine hyperperistalsis and dysperistalsis as dysfunctions of the mechanism of rapid sperm transport in patients with endometriosis and infertility,” Human reproduction, 11(7), pp. 1542–1551.

[34] Kunz, G. and Leyendecker, G., 2002, “Uterine peristaltic activity during the menstrual cycle: characterization, regulation, function and dysfunction,” Reproductive biomedicine online, 4, pp. 5–9.

[35] Kissler, S., Hamscho, N., Zangos, S., Wiegratz, I., Schlichter, S., Menzel, C., Doebert, N., Gruenwald, F., Vogl, T., Gaetje, R., et al., 2006, “Uterotubal transport disorder in adenomyosis and endometriosis—a cause for infertility,” BJOG: An International Journal of Obstetrics & Gynaecology, 113(8), pp. 902–908.

